# A simple method for comparing peripheral and central color vision by means of two smartphones

**DOI:** 10.1101/2021.01.12.426150

**Authors:** Galina I. Rozhkova, Alexander V. Belokopytov, Maria A. Gracheva, Egor I. Ershov, Petr P. Nikolaev

## Abstract

Information on peripheral color perception is far from being sufficient since it was predominantly obtained using small stimuli, limited ranges of eccentricities, and sophisticated experimental conditions. Our purpose was to consider a possibility of facilitating technical realization of the classical method of asymmetric color matching (ACM) developed by Moreland and Cruz (1959) for assessing appearance of color stimuli in the peripheral visual field (VF). We adopted the ACM method by employing two smartphones to implement matching procedure at various eccentricities. Although smartphones were successfully employed in vision studies, we are aware that some photometric parameters of smartphone displays are not sufficiently precise to ensure accurate color matching in foveal vision; moreover, certain technical characteristics of commercially available devices are variable. In the present study we provide evidence that, despite these shortages, smartphones can be applied for general and wide investigations of the peripheral vision. In our experiments, the smartphones were mounted on a mechanical perimeter to simultaneously present colored stimuli foveally and peripherally. Trying to reduce essential discomfort and fatigue experienced by most observers in peripheral vision studies, we did not apply bite bars, pupil dilatation, and Maxwellian view. The ACM measurements were performed without prior training of observers and in a wide range of eccentricities, varying between 0 and 95°. Color appearance was measured in the HSV color space coordinates as a function of eccentricity and stimulus luminance. We demonstrate that our easy-to-conduct method provides a reliable means to estimate color appearance in the peripheral vision and to assess inter-individual differences.

## Introduction

In natural viewing conditions, the human visual system integrates input signals from central and peripheral retina creating a single perceived image of the whole visual field (VF).

It is well known that the sensory surface that receives the optical inputs from all parts of the VF – the retina – has a very heterogeneous structure. This implies that when visual stimuli fall onto different spatial locations in the retina the same visual task has to be solved by recruiting different neuronal “units” and mechanisms. Such a situation stimulated comparative studies of the visual processes at the center and periphery of the VF, in particular, concerning color perception.

However, it does not mean that the characteristics of the central and peripheral color vision have been studied equally well since investigation of the peripheral vision is much more difficult (Moreland, 1972; Abramov & Gordon, 1977; Neitz, 1990; Rozhkova et al., 2019). In everyday life, one’s attention is usually directed to the central area of the visual field, leading to preferential processing of the central visual stimuli while the experiments aimed at investigation of the peripheral vision require changing the habitual viewing mode and drawing attention to the peripheral stimuli. This explains why the majority of reliable studies of the peripheral vision were performed after special training of participants.

By now, the number of reliable studies of the peripheral color vision is rather small and does not yet allow drawing sufficiently complete picture of it. This implicates the necessity of new findings and, also, prompts development of novel experimental procedures. Such demand is increasing in view of recent discovery of new light-sensitive elements in the human retina, melanopsin-expressing ganglion cells (ipRGCs) that are considered to contribute to color perception, too.

One of the traditional methods for assessment of the peripheral color vision is an asymmetric color matching (ACM) proposed more than 60 years ago (Moreland & Cruz, 1959) and widely used in many investigations up to now (Stabell & Stabell, 1976, 1979, 1982a, 1984, 1996; McKeefry et al., 2007; Hansen et al., 2009; Murray et al., 2012; Opper et al., 2014; etc.). A classical color matching paradigm (e.g., realized in the Wright colorimeter) implies perceptual equalization of the two colors presented on the two halves of the colorimeter screen by means of proper adjusting the reference light composed of the three monochromatic light beams, the colorimeter primaries. In the Moreland and Cruz ACM procedure, a visual match is accomplished between one stimulus imaged on the central part of the retina, and another stimulus projected onto the peripheral part of the retina.

The two matching procedures are based on the assumption that the human color vision is trichromatic. This is shown to be at least true of the foveal color vision under daylight (photopic) conditions when color perception is determined by inputs of the three types of cone photoreceptors. However, in a general case, one needs to take into account other photosensitive retinal elements partaking in color vision. By now it is established that, at different light conditions and for different regions of the VF, the following retinal cells selectively sensitive to different wavelengths contribute to human color perception: three types of cones determining daytime color vision and contributing to the twilight vision (short-wave S-cones, with peak sensitivity at ~430 nm; middle-wave M-cones with peak sensitivity at ~530 nm; and long-wave L-cones with peak sensitivity at ~560 nm); rods, the receptors much more sensitive to light than cones (with peak sensitivity between S- and M-cones at ~520 nm) enabling night vision and contributing to twilight vision; and intrinsically photosensitive retinal ganglion cells (ipRGCs) containing melanopsin with peak sensitivity between rods and L-cones (at about 490 nm). The number of publications on the ipRGSs and their contribution to visual functions is rapidly growing (Berson et al., 2002; Hattar et al., 2002; Dacey et al., 2005; Graham, 2014; Cao & Barrionuevo, 2015; Baraas & Zele, 2016; Hannibal et al., 2017; Zele et al., 2018; Schroeder et al., 2018; Allen et al., 2019; Spitschan, 2019; Yamakawa et al., 2019, etc.).

Figure 1A presents information on spectral sensitivity of these retinal elements (gray curves); superimposed are the monochromatic primaries used in the classical Wright colorimeter (color bars).

**Figure 1.**
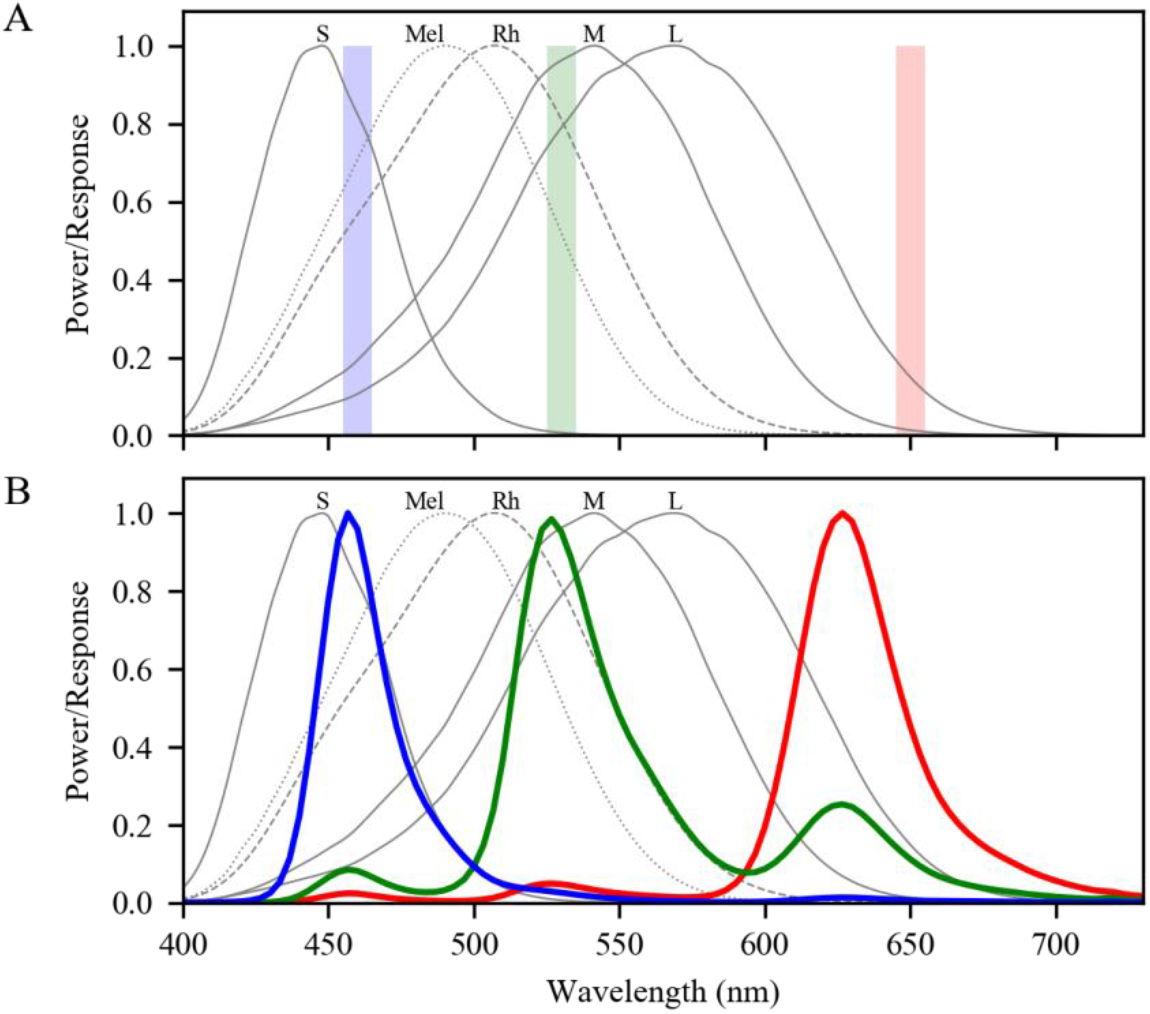
(A) The spectral sensitivity curves of rods (Rh), cones (S, M, L) and melanopsin ganglion cells (Mel) in the human retina according to the data from (Commission Internationale de l’Eclairage [CIE], 2018). The three colored bars indicate the Wright primaries commonly used in colorimetric experiments – monochromatic beams of 650, 530 and 460 nm (with each of 10 nm width). (B) Spectral characteristics of the smartphone (Samsung Galaxy S8) RGB primaries (colored curves) used in the present experiments as proxies of the Wright primaries and superimposed on the spectral characteristics of the color sensitive elements in the human retina.

It is well known that distribution of the light-sensitive cells over the human retina is very inhomogeneous (Østerberg, 1935; Curcio et al., 1987, 1990) as illustrated by Figure 2. Cone density is maximal at the retinal center (fovea) and decreases with eccentricity very steeply up to 3° (point 1 on the corresponding curve in Figure 2); in the range of 3-10° (between points 1 and 2 in Figure 2), the decrease is less; at the interval 10-80° (between points 2 and 3), the decrease is very slow, and at the retinal edge (point 4), near *ora serrata*, there is a significant increase in cone density known as cone-enriched rim (Williams, 1991; Mollon et al., 1998; To et al., 2011). In comparison, rod density has its peak at mid-periphery of the retina. Moreover, rod density as a function of eccentricity varies substantially for different radial directions. For instance, in the nasal part of the horizontal meridian, the peak rod density is confined to the range of 10-20° (over the 120*10^3^/mm^2^, indicated in Figure 2) compared to the 15-40° range in the temporal part of this meridian.

**Figure 2.**
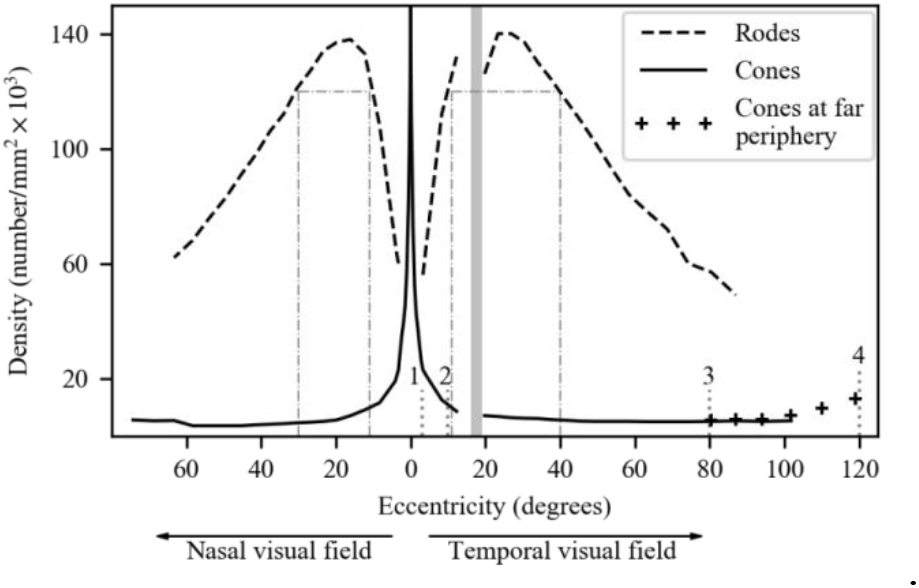
Inhomogeneous distribution of rods and cones in the human retina. Schematic illustration (based on Østerberg, 1935; Curcio, 1990). The points 1, 2, 3, and 4 correspond to the ends of the intervals with different cone density gradients. Dash-dotted vertical lines indicate widths of rod peak zones in the nasal VF (temporal retina) and temporal VF (nasal retina) along the horizontal meridian; gray bar indicates temporal VF eccentricity of the blind spot.

Currently the data on the distribution of the ipRGCs in the human retina are very preliminary. One could only be sure that the ipRGCs are absent in the central 2° of the retina since, in this area, all ganglion cells is moved aside to the edges of the fovea.

Till our days, many ophthalmologists and some visual scientists share the last century’s notion that, with increasing eccentricity, there is a transition from normal trichromatic vision to dichromatic vision, and utterly to achromatic vision (color blindness) at the far periphery. Sixty years ago, such a view was clearly formulated by Moreland and Cruz whose paper (Moreland & Cruz, 1959) influenced the peripheral vision investigations for many years. Their paper presented results of estimation of appearance of the peripheral colors by using, for comparison, a foveal mixture field. The authors wrote: “The results indicate a progressive deterioration in colour perception with distance from the fovea: tending, under the conditions of the experiment, to dichromatism at 25°-30° and to monochromatism at 40°-50°” (Moreland & Cruz, 1959, p. 117).

The data that justified their conjecture were obtained for two observers (the authors) under dark adaptation, monocular (right eye) viewing, and the observer’s head stabilized by a dental plate mounting. The test and comparison stimuli (80min x 40min) were presented simultaneously as pulses of 0.5 s duration, every 2 s. The position of the test stimulus was varied from 10 to 30° in the lower vertical meridian and from 15 to 50° in the nasal horizontal meridian. Although the authors explicitly indicated that their conclusions were based on the data obtained “under the conditions of the experiment” (Moreland & Cruz, 1959, p. 117), their view of the degraded peripheral color vision was often unlawfully generalized by successors in this field of research.

However, numerous post-1959 studies demonstrated that the original view on the peripherally perceived colors required profound revision, whereby convincing evidence was provided that, at the VF periphery, in their appearance the colors are comparable to the same as the foveal colors. if the peripheral stimuli are sufficiently large and bright (Abramov & Gordon, 1977; Gordon & Abramov, 1977; Smith & Pokorny, 1977; van Esch et al., 1984) and move across the retina with a proper velocity (Rozhkova & Yarbus, 1974; Yarbus & Rozhkova, 1978). For instance, van Esch et al. (1984, p. 443) wrote: “If field-size scaling according to the eccentricity-dependent cone density, the cortical magnification factor, or the reciprocal of the interganglion cell distance is applied, then wavelength-discrimination performance from 8 degrees to 80 degrees eccentricity is roughly the same.”

By now, the majority of the studies on peripheral color vision report data obtained at rather limited and highly-specified conditions (Maxwellian view; adaptation procedures; limited range of eccentricities, stimulus size, luminance, temporal parameters, ambient illumination levels, etc.). It is therefore hardly of surprise that under different experimental conditions, quite different phenomena were recorder and, thus, different (even opposite) conclusions were arrived at.

For instance, one might compare the following conclusions drawn by several authors about perceived saturation of the peripheral stimuli:

1. “The quality of color vision in the periphery depends crucially on stimulus size. If the stimulus is sufficiently large, subjects see a full range of well saturated hues.” (Gordon & Abramov, 1977, pp. 205-206);
2. “In agreement with previous studies the results show that the saturation of the different colours is reduced when the monochromatic stimuli are moved from the fovea toward the periphery” (Stabell & Stabell, 1976, p. 1100);
3. “Foveal and peripheral hue-scaling data were obtained for a 1° foveal stimulus and a 3° stimulus presented at 10° retinal eccentricity under both bleach (reducing rod input) and no-bleach (permitting rod input) conditions. … Peripheral stimuli appeared more saturated than foveal stimuli (i.e., supersaturated), especially in the green–yellow region. This effect was particularly pronounced for the peripheral bleach condition” (Opper et al., 2014, p. A148).
4. “The task for the observers was to match a peripheral 3°-spot at 18°-eccentricity in the nasal VF with a parafoveal 1°-probe spot (1°-eccentricity, nasal VF) in hue and saturation. For both males and females, there is greater saturation loss in the green region with males exhibiting higher loss. Females show a slightly greater saturation loss in the red region of the color space” (Murray et al., 2012, p. 3).

The above quotations make the available data seem like a collection of disjointed fragments of information in a puzzle with too many tiles absent, which precludes working out a complete view of the peripheral color vision.

From the viewpoint of ecological validity, the main hindrance in understanding of the peripheral color vision phenomena is lack of data obtained under natural viewing conditions and by accessible methods. The traditional lab-based investigations of the peripheral color vision have their limitations, such as the technically elaborate Maxwellian view; expensive equipment required for colorimetric measurements; long lasting and complex adaptation procedures, etc., i.e. the factors that limits research potential to acquire a richer information on the topic.

We consider that development of an experimental technique that is accessible and affordable for wide application in research actual since it opens new avenues to enhance investigation of the peripheral vision. Building upon the requirements of accessibility and broad applicability, we developed an easy-to-handle setup with two off-the-shelf smartphones for conducting a pilot study to compare and match in appearance colors presented centrally and in the periphery of the VF. Although we are aware that off-the-shelf smartphones are devices that not suitable for performing precise measurements, we reckoned that they are convenient for a rapid assessment of certain characteristics of matched colors and for demonstration of certain phenomena of peripheral color vision.

In our experiments, we employed smartphones to implement the ACM procedure. Figure 1B shows spectral characteristics of the smartphone primaries which are proxies of the three monochromatic beams (cf. Figure 1A) used as the primaries in the classical Wright colorimeter. As one can see, the bandwidths of the smartphone primaries are much wider and the spectrum of the green primary has significant side peaks. However, from a theoretical viewpoint, such particularities are not crucial in colorimetry (the first law of Grassmann only requires linear independence of the primaries (MacAdam, 1970; Wyszecki & Stiles, 1982)).

The paper contains two main parts. The Methods section provides information on technical and photometric characteristics of the employed smartphone that are essential for the present study, details of the experimental setup, and the ACM procedure.

The Results section presents our findings on the appearance of color stimuli presented peripherally as matched to those presented foveally that justify the proposed technical elaboration and the method. The section contains novel data and provides results that confirm some earlier outcomes in studies that employed more sophisticated lab equipment and procedures, and trained observers.

Some of our experiments were carried out for clarifying certain technical issues – the smartphone temperature, temporal properties of the stimuli, benign differences in spectral characteristics of the test and reference devices. Outcomes of these auxiliary experiments are presented in Supplement.

## Methods

### The experimental setup

Our main experiments were carried out using a pair of smartphones with supposedly identical characteristics and a modified commercial mechanical perimeter PNR2 (analogous to the Foerster perimeter) with an arc radius of 33 cm. The basic setup is shown in Figure 3A. The smartphone for the peripheral test stimulus presentation was attached to a platform that could be moved over the elongated perimetric arc to vary eccentricity of test images. The second smartphone was fixed at the center of the perimetric arc (at zero eccentricity) for displaying the reference color images.

**Figure 3.**
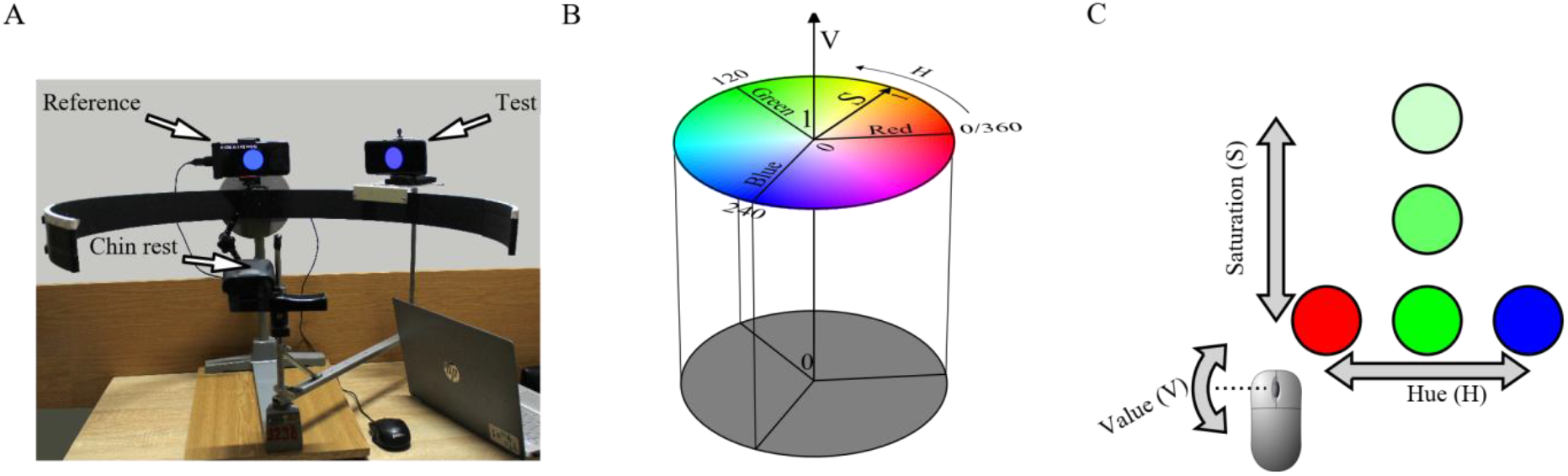
(A) Experimental setup for color matching: Photo of the perimeter with two smartphones for displaying test and reference stimuli. (B) Schematic representation of the HSV color coordinate system used in this investigation. (C) Scheme of adjusting reference image color by means of manipulations with mouse. The peripheral smartphone was connected to a notebook for test image generation and setting its parameters. Image parameters on the central smartphone could be varied by means of manipulations with a mouse connected to this smartphone. Quantitative characteristics of the displayed color were expressed in HSV color coordinate system. (Some reasons in favor of using HSV color space in our experiments are given below; a schematic representation of HSV color space is shown in Figure 3B.) The observers had to match the central reference stimulus to the peripheral test stimulus in hue (H), saturation (S), and brightness (V). This matching was achieved by moving the mouse in different directions (to vary H and S) and rotating the mouse scroll wheel (to vary V) (Figure 3C). The software for color image generation on the smartphones was developed using JavaScript (the source code is accessible on https://github.com/abelokopytov/spot).

### The choice of the color coordinate system

For estimation of color appearance, we preferred to use well known HSV color coordinate system elaborated in 70s of the previous century (Joblove & Greenberg, 1978; Smith, 1978; Wyszecki & Stiles, 1982) although, since those times, many other color coordinate systems were developed for various visual tasks and applications.

For our investigation, the type of color coordinate system was not very crucial. To make the color choice easier and faster and, hence, to reduce the duration of the matching experiments, the following requirements seemed to be reasonable:

1. The system should be designed for image visualization on displays and should meet the characteristics of smartphones providing representation of all possible colors (smartphone color gamut).
2. The system should be easy to understand by naïve observers, i.e. it should be correlated with visual perceptual qualities of color encountered in common use – hue, saturation, and brightness/lightness.
3. It is desirable to separate the control of image chromaticity from image intensity and to provide simple adjustment of color images by mouse (e.g. movements of the mouse over the table may determine changes of chromaticity while turning the mouse wheel – changes of brightness).
4. The system should be suitable for the meaningful association of different mouse movements with adjustment of different chromaticity parameters (e.g. left-right directions of movement – with hue changes while forward-backward directions – with saturation changes).

HSV color coordinate system belongs to RGB class. In general, color systems of RGB class are more suitable than others for the images generated on displays. This class includes about a dozen versions. However, most of them don’t fit at least one of our requirements. In particular, the systems CMY и CMYK don’t meet the point (1) since they are developed for printed images. The requirement (3) excludes widespread systems RGB, YUV, and ICtCp since, in these systems, each of the three free parameters influence stimulus chromaticity. By taking into account the requirement (4), the list of candidates shortens to HSV and several rarely used color coordinate systems. In such a situation, it seems rational to choose HSV because it is familiar to many investigators and describes the colors in terms (hue, saturation, and value=intensity) easily accepted by naïve observers.

The cylindrical scheme of HSV color space presented in Figure 3B shows the top disk corresponding to maximum intensity colors and the bottom disk corresponding to blackness / darkness. The cylindrical surface between the two circular contours contains the maximum-saturated colors for all intensities and for all hues presented in the circularized chromaticity diagram.

### General information on the employed smartphone models

In a majority of color matching experiments, the two smartphones Samsung Galaxy S8 with AMOLED screen were employed. At the beginning of the investigation, another pair of smartphones was also used: Samsung Galaxy S6 with an AMOLED screen (see Table 1).

**Table 1.** Details of the smartphone models employed in the present study.

| Model | Display Type | Screen Resolution (Pixels) | Pixels Per Inch | Released |
| --- | --- | --- | --- | --- |
| Galaxy S6 | Super AMOLED | 2560x1440 | 576 | 2015 |
| Galaxy S8 | Super AMOLED | 2560x1440 | 568 | 2017 |

We have chosen the smartphones with OLED screens (not LCD) because they have excellent contrast ratios and can provide high brightness and rich gamut.

### Spectral characteristics of the smartphone primaries

It is apparent that, ideally, for accurate matching central and peripheral images, the test and reference smartphones should have identical spectral characteristics of their R, G, and B primaries. Figure 4 shows the two sets of RGB spectra for our test and reference smartphones of the two Samsung — Galaxy S6 and Galaxy S8. All spectral characteristics were obtained with X-Rite Eye One spectrophotometer. The measurements were carried out taking into account the period necessary for stabilization of the smartphone characteristics after its switching on. The duration of this period was assessed in advance and did not exceeded 5 min.

**Figure 4.**
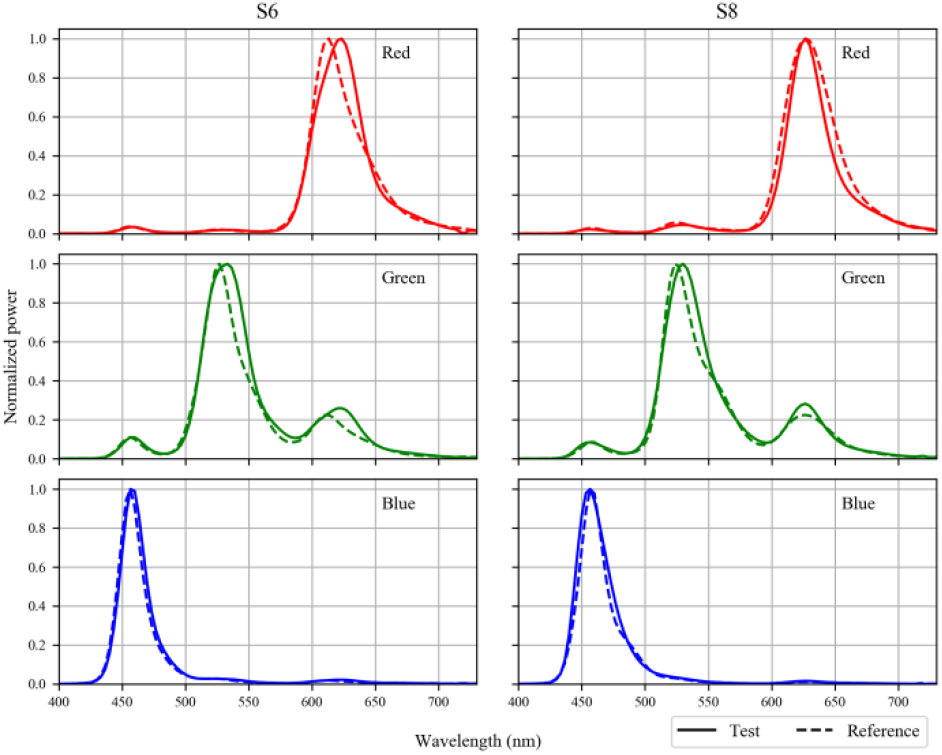
Spectral characteristics of the R, G, and B primaries for the test-reference pairs of smartphones Samsung Galaxy S6 and S8 used in the experiments. Solid lines correspond to the central reference smartphone, dashed lines – to the peripheral test smartphone.

As one could see from Figure 4, there were small differences both between the smartphone models (S6 and S8) and between the two devices (test and reference) of the same model. All corresponding curves appeared to be similar but not quite identical. Comparing the differences between spectral characteristics of the primaries in the test-reference pairs of the models S6 and S8, it is easy to reveal that S8 has some advantages since the conformity of the curves in the pairs of its red and green primaries is better (at least, as concerns the proximity of their peaks).

Unfortunately, the technical imperfectness could not be eliminated. Therefore, it was important to evaluate the influence of the described differences on the results of ACM. For this purpose, we performed the same series of measurements interchanging test and reference smartphones. The data obtained in such comparative experiments are presented in the Appendix.

### Calibration of the smartphones before the color matching

Interpretation of the ACM experimental data requires some supplementary information about the smartphone calibration and specific details of the matching procedures. Calibration procedures included: (1) setting of each smartphone luminance at the chosen level for white stimulus; (2) measuring physical luminance L for the three color primaries displayed on the test and reference smartphones at different luminance levels, L_sm_, expressed in 8-bit 256 RGB values; (3) finding the relationship between the brightness parameter V (varying from 0 to 1) of the reference image and the test image luminance L_sm_ (in 8-bit 256 RGB values).

All photometric procedures were performed with Gossen MAVO-MONITOR luminance meter. The luminance of each smartphone “white” (R=G=B=255) was set using Android application “Brightness adjuster” by K. Shimokura at the level of 400 cd/m^2^ under photometric control. Such level was chosen in order to have a possibility to present sufficiently bright test stimuli within the zone of comfort for human visual perception but not too close to the very highest limit (500 cd/m^2^) of the devices employed. The adjusting devices on the basis of “white” is one of the traditional procedures in the experiments with color perception aimed at investigation of vision in natural conditions while, in threshold investigations, quite a different setting might be required.

The relationships between L_sm_ in 8-bit RGB values and the photometric data of L measurements are shown in Figure 5. The curves corresponding to the test and reference smartphones (solid and dashed lines) appeared to be practically identical.

**Figure 5.**
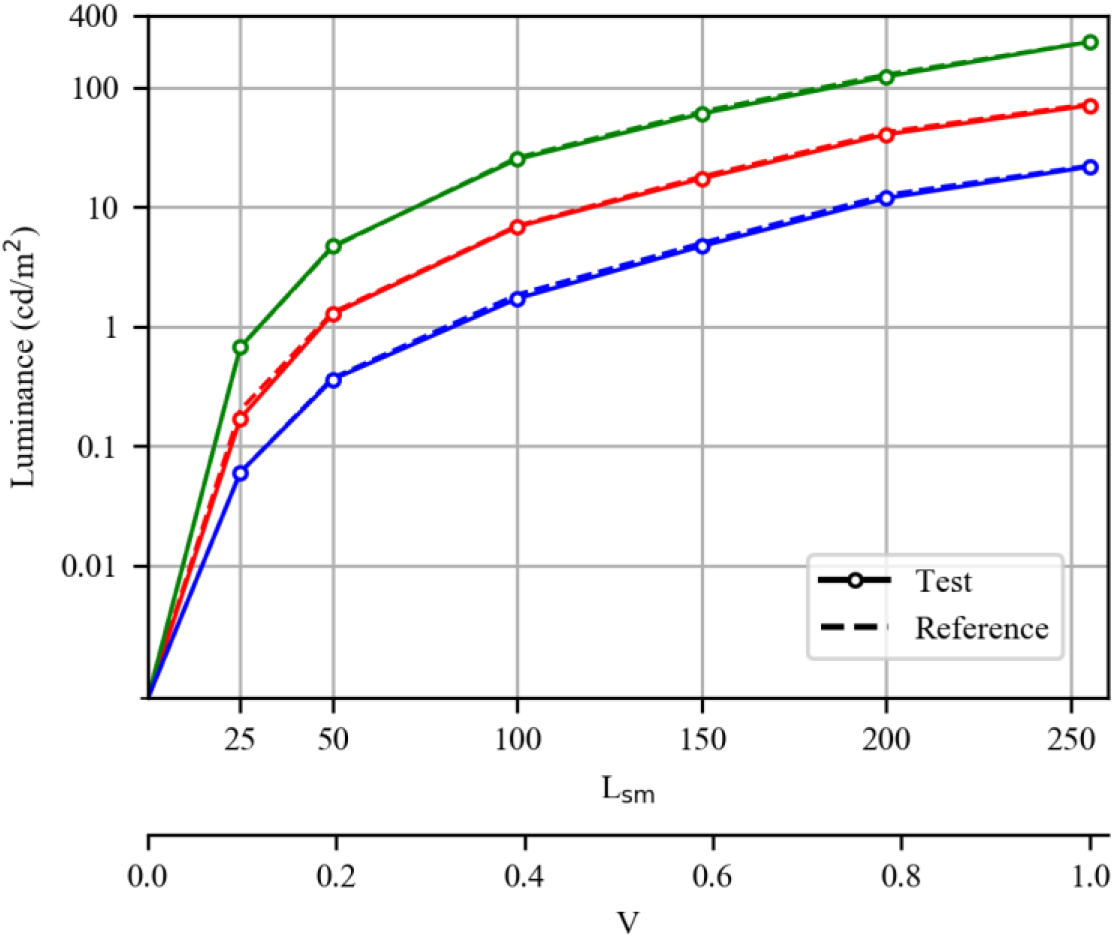
Relationships between L_sm_ (in 8-bit RGB values), value V (in HSV color space) and photometric luminance L for the 3 primaries of the test and reference smartphones (solid and dashed lines, respectively).

To estimate the smartphone’s display gamma, we used manual fitting and got gamma=2.4. Usually, in colorimetric measurements with computer displays, one sets gamma to 1, getting linear dependence of luminance on pixel value (ranging from 0 to 255 in 8-bit presentation of colors). But still, there are no easy ways to change display gamma in android v.9 used in our Samsung Galaxy S8 smartphones.

Some additional information characterizing the smartphone properties substantial for our experiments is given in Appendix: effect of the temperature on the luminance of each primary; effect of the reference stimulus temporal mode (flickering vs stationary) on the results of matching; effect of interchanging the test and reference smartphones.

### Participants

Participants were eight volunteers (3 females), aged 19-46 years. According to the results of a standard clinical examination (assessment of refraction, accommodation, visual resolution, and color discrimination), all participants had normal visual acuity and normal trichromatic color vision. As concerned color matching measurements, three participants were experienced while the other five were naïve observers. Our investigation included two stages: (1) a preliminary stage consisting of short experimental sessions aimed at assessing suitable stimulus parameters, refining the experimental protocol, and obtaining initial tentative results; (2) the main stage aimed at collecting data sufficient for thorough analysis and reliable conclusions. All eight observers participated in the preliminary stage providing an opportunity to develop reasonable experimental protocols and to assess inter-subject variability of responses. However, only half of the observers could participate in the main experiments that were rather tiresome and whose duration was substantial.

The study was carried out in accordance with the Declaration of Helsinki (1975). Prior to testing, written informed consent was obtained from each participant. The experimental procedure was approved by the Institute for Information Transmission Problems ethics board.

### Stimuli

The peripheral test images were uniform color disks of 4 cm (about 7°) in diameter turning on/off each 1.5 s. The observation distance was 33 cm (the radius of the perimetric arc). The size of the test images was large enough to make them well seen at the periphery. Low-frequency flicker was used to provide comfortable conditions for prolonged observation of the test images at far periphery taking into consideration well known Troxler effect – fading of stationary peripheral stimuli in conditions of gaze fixation on the center of the VF (Troxler, 1804). The duration of “on” and “off” phases were chosen so as (1) to prevent such fading, (2) to provide suitable temporal conditions for matching procedure, and (3) to avoid the development of noticeable afterimages arising after the stimulus offset in the cases of high stimulus luminosity and long duration. Performing several preliminary experiments with “on” and “off” phase durations of 1.0, 1.5, and 2.0 s for comparison, we became convinced that the results were similar but the presentation mode 1.5/1.5 s seemed to be somewhat more comfortable for our observers.

The color of each test stimuli corresponded to one or another of the smartphone primaries – R, G, or B (their spectral curves were shown in Figure 1B and Figure 4). The luminance of the test stimuli (L_sm_) was set in 8-bit 256 RGB values: 25≤R≤255, G=0, B=0 for the red stimuli; R=0, 25≤G≤255, B=0 for the green ones; R=0, G=0, 25≤B≤255 for the blue ones. In the course of each experimental session, six levels of L_sm_ were used for red, green, and blue images: 25; 50; 100, 150, 200, and 255. The perceived color of the peripheral test image was estimated by proper adjusting the central reference image and was expressed in HSV color coordinate system. In this notation, all the test stimuli should ideally have the following perceived hue values H in an angular dimension: H(R)=0/360; H(G)=120; H(B)=240. The accepted range for the intensity parameter V (the value characterizing perceived brightness) is from 0 to 1.0. As concerned the perceived saturation S, the possible range is also from 0 to 1; ideally, for all our test images, this parameter should always be maximal (S=1), since each test image was a pure smartphone primary. At the same time, the spectral curves shown in Figure 2 indicate that green stimuli could be perceived as somewhat less saturated than red and blue ones because of the larger additional lateral peaks.

The reference images were of the same size as the test ones, however, in order to ease matching, the reference images were not turning on/off since these images had to be permanently adjusted.

The physical levels of the image luminance (L in cd/m^2^) corresponding to each L_sm_ value could be determined using the calibration curves presented in Figure 5.

The following eccentricities along the horizontal meridian in the temporal half of the VF were chosen for detailed measurements: 0, 25, 40, 60, 80, and 95°. Unequal intervals at the beginning and at the end of the eccentricity range were used in order not to be near the regions corresponding to the blind spot and the margin of the retina, *ora serrata*.

### Procedure

The stimuli were presented in a dimly lit room with the illumination of 5 lux. At the beginning of each measurement session, the observers were given 5 min to adapt to this level of ambient light and to familiarize themselves with the equipment. The experiments were carried out in conditions of monocular viewing and keeping the gaze on the central reference image. In the course of measurements, the participant was seated in such a way that his/her tested right eye axis was directed to the reference image at the center of the perimetric arc. The left eye was covered with an opaque patch. The observer’s head was supported by a chin rest. The peripheral test stimulus displayed on the smartphone at variable eccentricity and the reference image displayed on the second smartphone located centrally were observed simultaneously. Since the reference stimulus had the same relatively large size as the test one (7°), it would be not quite correct to use the term “foveal fixation”. According to a widely accepted view, the diameter of the foveal area is about 5° (unfortunately, there are no reliable data on inter-subject variability of this parameter), therefore, it is a more proper way to say that our observers were examined under conditions of central fixation and that the variation of the eccentricity of the peripheral test stimulus was to a larger extent than under conditions of using a fixation point at the very center of the VF.

The participant’s task was to make the appearance of the central reference image as similar as possible to the appearance of the peripheral test stimulus adjusting perceived hue (H), saturation (S), and brightness (V) of the reference image. The parameters of the reference image could be easily controlled by participants with a mouse connected to the reference smartphone. In order to change the hue and saturation of the reference image, the participant has to move the mouse left-right and forward-backward, respectively; brightness can be adjusted by means of turning the wheel. The time given for matching was not limited. Each trial could take several seconds or one-two tens of seconds depending on the participant’s capabilities and test image parameters. Once a satisfactory match was reached, the participant pressed the button of the mouse to save the result. The recorded HSV values were displayed on the reference smartphone after each completed trial and disappeared before the next trial.

Most participants were tested during 2-5 days since the final standard protocol of our main experiments included too many measurements to be completed in one working day. Participants had to perform matching for 3 primaries (R, G, B) and obtain three estimates (H, S, and V) at each of 6 luminance levels (25, 50, 100, 150, 200, and 255 in RGB-scale) for the test stimuli presented at six eccentricities (0°, 25°, 40°, 60°, 80°, and 95° on the horizontal meridian of the temporal VF). All measurements were repeated 5 times. The final dataset for each participant contained at least 3 x 3 x 6 x 6 x 5=1620 values.

## Results

Since our main purpose was to consider the possibilities and the specifics of the proposed technique, we focused our attention on encompassing the whole range of eccentricities and obtaining comprehensive data on general features of the peripheral color vision.

Presenting color test stimuli at the periphery (temporal VF, 25-95°), we have found that, in the whole range of the eccentricities used, the perceived peripheral images could be clear and vivid. However, it is necessary to mention an important peculiarity of the colors perceived at the periphery: though, in most cases, the observes could choose the central image most similar to the peripheral test image, the task of setting a completely identical central image on the reference smartphone was not always feasible though the response time was not limited. For this reason, each participant was instructed to do his/her best but not to be too anxious if the desired perfect matching appeared to be unattainable.

From the description of our matching method, it is evident that, in each measurement, all the three quantitative estimates of the perceived peripheral color – H, S, and V – were obtained simultaneously. These estimates were displayed at the reference smartphone just after the participant had chosen the most suitable central image and pressed the mouse button. However, in order to simplify the description and interpretation of the experimental data, H, S, and V estimates will be considered separately.

### Perceived hue of the peripheral stimuli, H-estimates

The data characterizing the dependence of H-estimate on the test image luminance L_sm_ and eccentricity are shown in Figure 6. This figure contains H-estimates obtained in 4 participants for 5 levels of the smartphone luminance L_sm_ (25, 50, 100, 150, 200, and 255) and 6 eccentricities (0, 25, 40, 60, 80, and 95°). At the lowest luminance used in experiment, L_sm_ = 25, the participants were not always sure in their matches and we discard these measurements.

**Figure 6.**
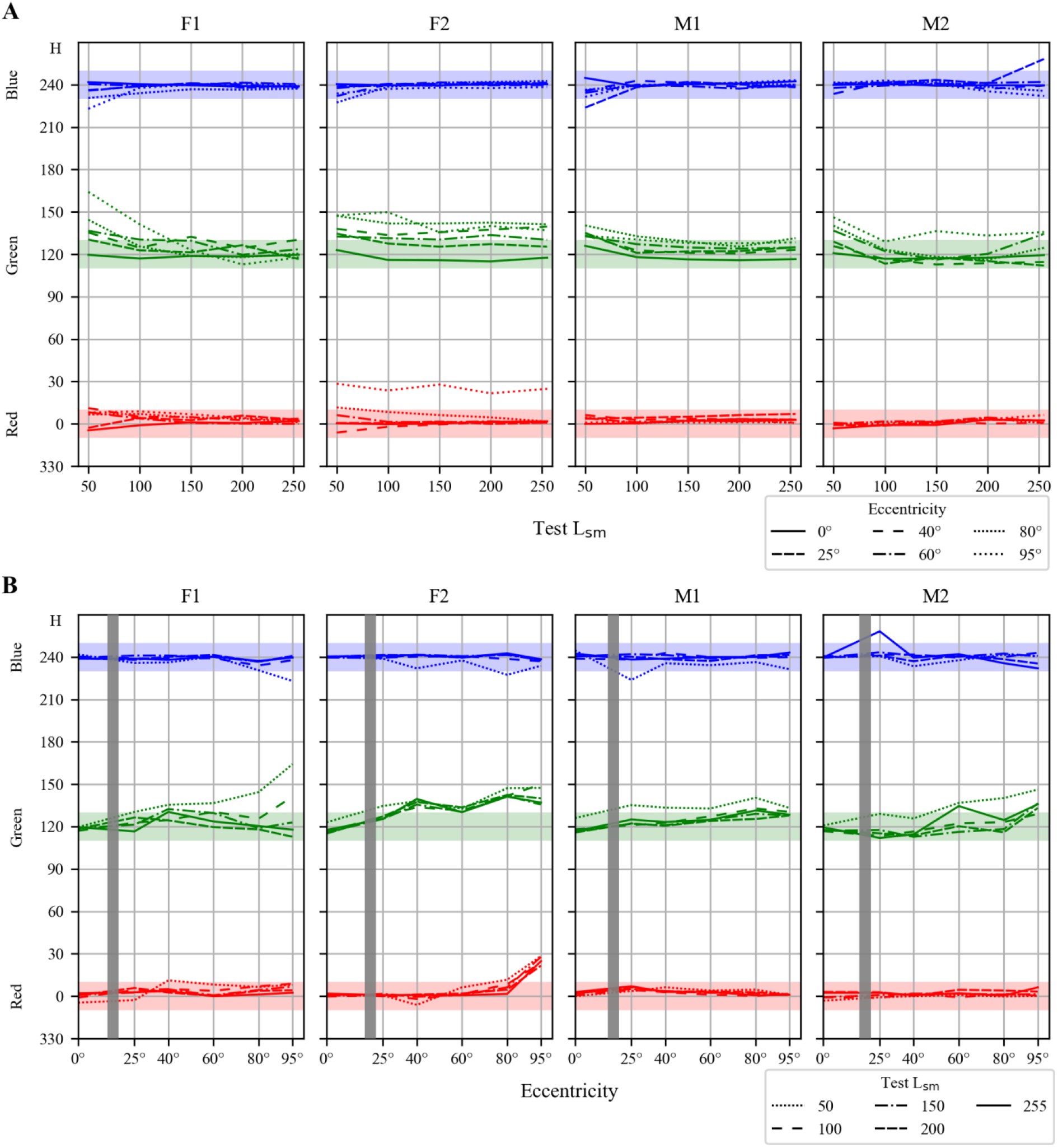
Perception of hue (H) at various luminance levels (L_sm_) and eccentricities of the peripheral test stimuli. Data for four participants, – two females (F1 and F2) and two males (M1 and M2) are presented to illustrate inter-individual differences. (A) – H as a function of L_sm_ at the varying eccentricity. (B) – H as a function of the eccentricity at varying L_sm_. Gray bars indicate the angles corresponding to the blind spot in the retina of each participant.

Figure 6A presents the curves showing the dependence of H on L_sm_ for 6 eccentricities. As one could conclude from this figure, in the case of the blue peripheral stimulis, the H-estimates could be close to the true value of the central ones (240) up to the largest eccentricity of 95° in all 4 participants and all individual lowest L_sm_ levels providing good similarity did not exceed 100.

In the case of the red stimuli, the peripheral H-estimates could be close to the central ones (0/360) up to the largest eccentricity of 95° in 3 participants (F1, M1, M2) from 4; like in the case of red stimuli, their individual lowest L_sm_ levels providing good similarity did not exceed 100. In the participant F2, at the eccentricity of 95°, the same red stimuli were perceived as the orange ones with H-estimates from 22 to 28, independently of L_sm_. In conditions of our experiments, this participant could only perceive the peripheral red stimuli as pure red disks up to the eccentricity of about 60°. It is worth to note that, in the participant F3 (not included in Figure 6), at the eccentricity of 95°, H-estimates were changing from H=110 (yellow-green) to H=0 (pure red) with increasing L_sm_.

In the case of the green test stimuli, at the eccentricity of 95°, H-estimates of the two participants (F2 and M2) did not approach the “true green” value (H=120) with increasing luminance even at the highest L_sm_ levels. In F2, the perceived colors remained somewhat bluish (130÷140) with increasing intensity even at the lesser eccentricity of 80°.

Certain features of the perceived color changes at the periphery of the VF are better seen in Figure 6B showing the dependence of H on eccentricity for 5 values of L_sm_. Many of the dotted lines corresponding to the lower levels of luminance, L_sm_=50, demonstrate significant deviations of the perceived hue at the periphery from the central one. In some cases these deviations were increasing with eccentricity: for instance, see H-estimates for green test images in subjects F1 and M2. However, in other cases, the curves were less monotonic, evidently because of the errors indicating an insufficient number of trials and/or significant influence of certain individual factors besides eccentricity.

### Saturation of the peripheral stimuli, S-estimates

The data on the saturation of the peripheral stimuli obtained in the described series of our experiments are presented in Figure 7.

**Figure 7.**
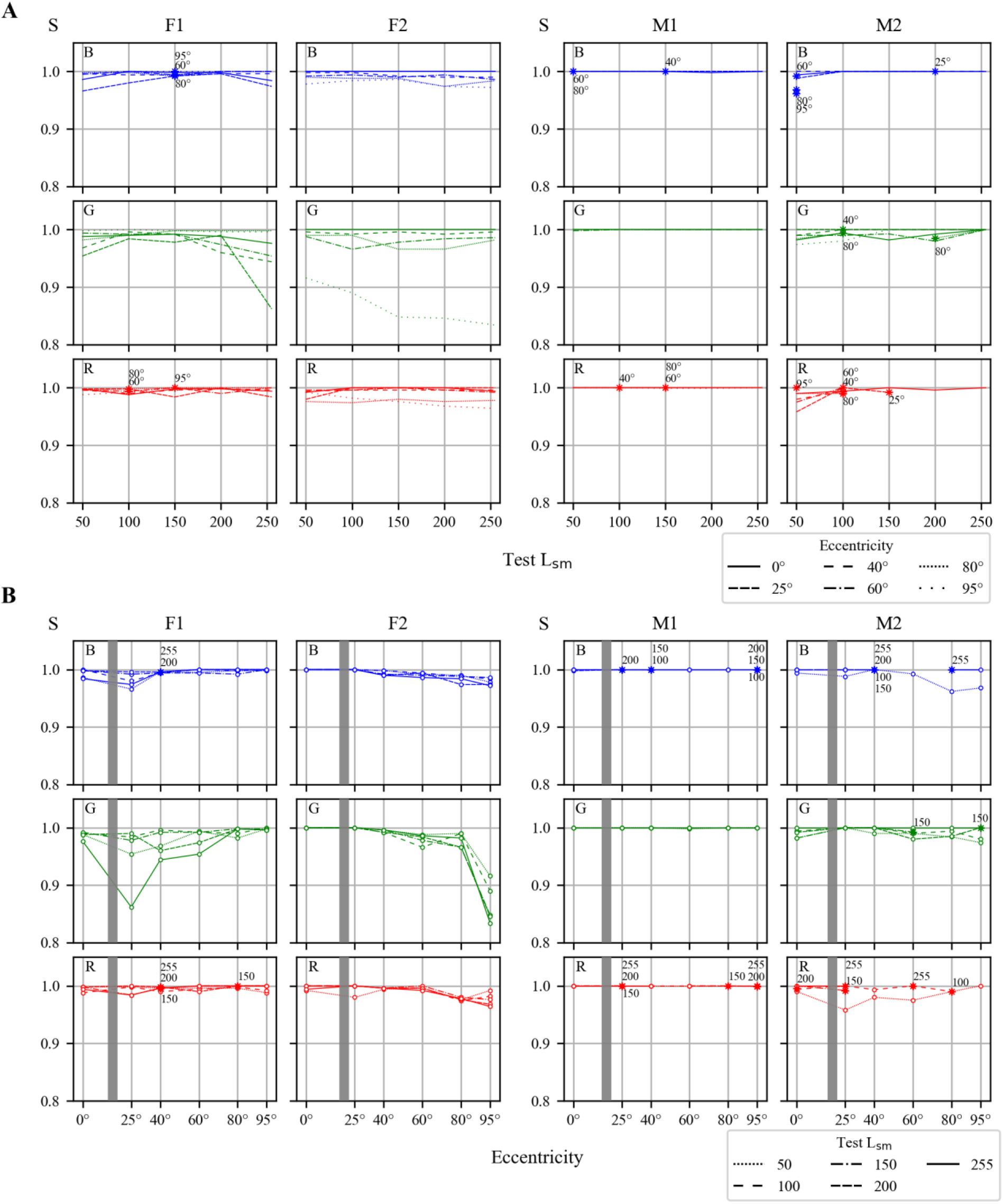
Saturation (S) as a function of stimulus luminance (L_sm_) and eccentricity. The data for four participants, two females (F1 and F2) and two males (M1 and M2). A) – S as a function of L_sm_ at different eccentricities: B) – S as a function of eccentricity at different L_sm_. Gray bars indicate the angles corresponding to the blind spot in the retina of each participant.

As was indicated in Methods, our color test stimuli always corresponded to the RGB primaries of the smartphones used in the experiments. Thus, the spectra of the stimuli were the same for all stimulus intensities and it was natural to expect that all estimates of saturation should be equal or close to the value S=1 prescribed to pure R, G, and B primaries, independently of their luminance.

In most cases, the data obtained for R- and B-images at lower values of L_sm_ (<150) and eccentricities (<60°) appeared to be close to this expectation: the preponderance of S-estimates was between 0.95 and 1. Because of such narrow range, many curves appeared to be very near to each other or even fused.

However, at far periphery, certain specific difficulties arose with the saturation of the bright test stimuli (L_sm_ >150). Such stimuli could look like more saturated than the central reference ones with S=1, thus making matching in S forbidden since, in the HSV color space, S>1 is inadmissible. In Figure 7, such cases are marked with stars (*) on corresponding curves. Since many curves are superimposed, appropriate eccentricities (Figure 7A) and L_sm_ levels (Figure 7B) are indicated near the curves.

The situation with G-images was more complicated, supposedly, because of the lateral peaks in the spectrum of the smartphone G-primary. On average, in the case of green stimuli, the correspondence between the peripheral and central S-estimates was worse than in the cases of Rand B-stimuli. At the same time, it’s noteworthy that this “shortage” of green stimuli was also well expressed in calibration experiments when both the test and the reference stimuli were perceived at the center of VF.

It was also noticed that, in some cases, when the participants perceived the peripheral image as “superior” to the central one with S=1 and V=1, they couldn’t separate contributions of saturation and brightness to this superiority. In such situations, the uncertain estimates “>1” could be prescribed to V, S, or both of them. One could suppose that our participants needed some training in the assessment of saturation. Though their oral responses evidenced against this assumption, we are going to check it in the near future.

### Brightness of the peripheral stimuli, V-estimates

The brightness (V-estimate) of the peripheral image was typically higher than that of the central one; the difference being maximal in the range of eccentricities 40-80°. The combined data of four participants in Figure 8 illustrate this finding showing the dependence of the brightness estimates V on the L_sm_ level at various eccentricities (A) and the dependence of the brightness estimates V on the eccentricity for different L_sm_ levels (B). The curves of Figure 8A evidence that, at the periphery, in contrast to the central region, the dependence of V on the L_sm_ is essentially nonlinear. This feature is better expressed around the eccentricity of 60°.

**Figure 8.**
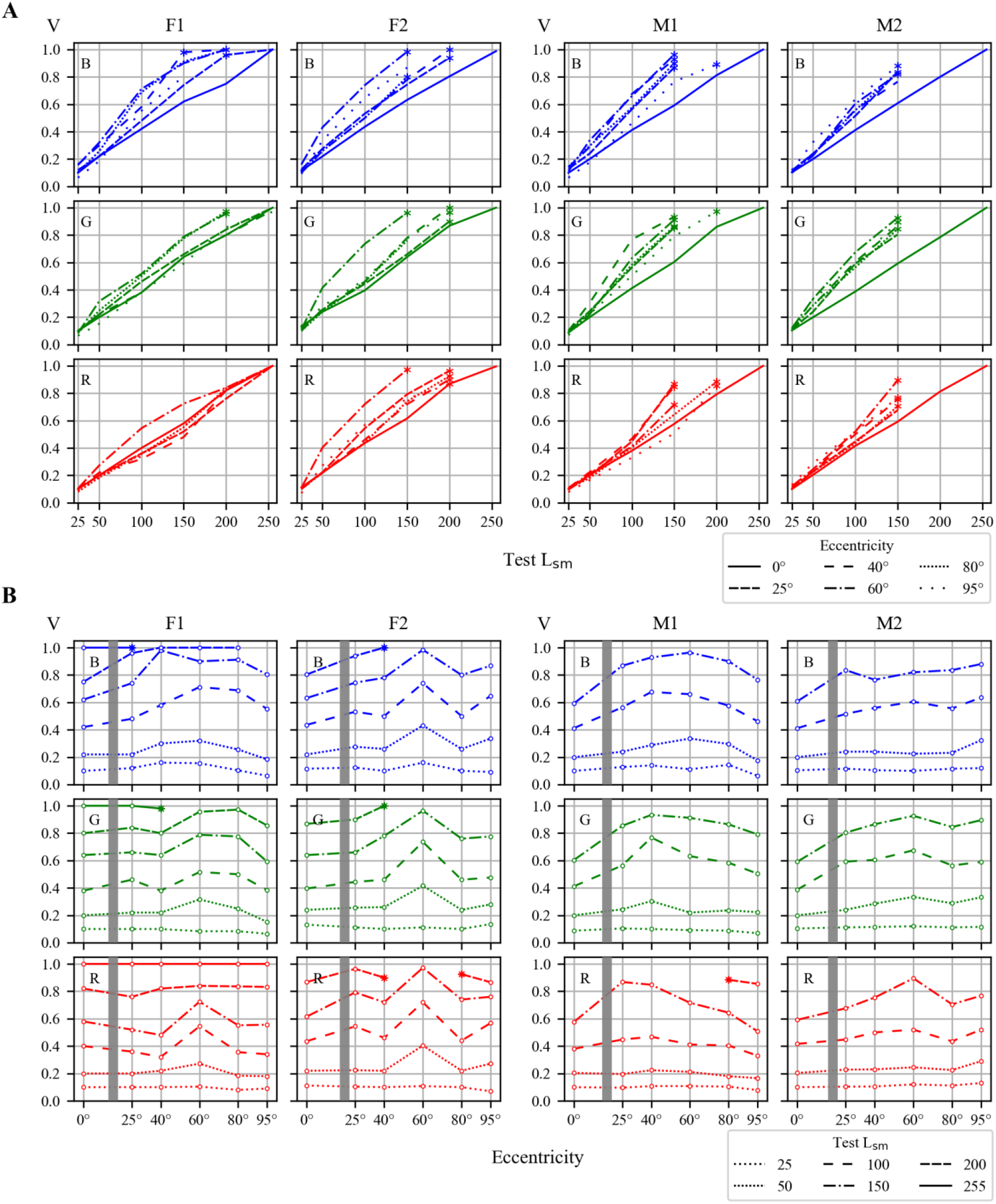
Brightness (V) as a function of stimulus luminance (L_sm_) and eccentricity. The data for four participants, two females (F1 and F2) and two males (M1 and M2). (A) – V as a function of L_sm_ at different eccentricities. (B) – V as a function of eccentricity at different L_sm_. Gray bars indicate the angles corresponding to the blind spot in the retina of each participant.

As one could conclude from Figure 8A, the brightness of the peripheral red, green, and blue images was higher around 60°, than at 0° for all participants and for all levels of L_sm_ exceeding 25. Notice that some curves in Figure 8 were marked with stars and were not completed. Like in the case of S-estimates, the stars correspond to the responses “>1” marking the L_sm_ levels (Figure 8A) or eccentricities (Figure 8B) over which the exact matching in V was impossible. In such cases, the limiting value of the reference central image, V=1, appeared to be “too small” for reflecting the brightness of the peripheral test image, and we couldn’t obtain any quantitative measure of its superiority over the central image.

Correspondingly, in Figure 8A, most curves obtained for the peripheral test images reached the brightness level V=1 at the L_sm_ values essentially lower than 255, the value which should theoretically correspond to V=1 according to the color perception model taken as a basis in HSV color coordinate system for trichromatic human central color vision. Moreover, in most cases, the data corresponding to the higher L_sm_ level couldn’t be presented in Figure 8 since the matching in V was impossible: the participant reported that the peripheral test image looked as more or even much more bright than the most bright central image that could be generated (corresponding to V=1).

The graphs of Figure 8B give a possibility for better understanding a general dependence of the brightness on eccentricity. More detailed analysis of Figure 8B reveals a significant inter-subject variability in the peripheral perception of brightness.

In participant F2, the data for R-, G-, and B-stimuli were similar to each other, the maximal V-values were obtained at the eccentricity of 60°. At the extreme periphery of 95°, most V-estimates were also obviously higher than the estimates of central images. In participant F1, the data for R-, G-, and B-images appeared to be essentially different; the positions of the largest V-values were varying in the range 40-80°. The difference between V-values at the extreme periphery and in the central region was small barring the cases of the two upper curves for the blue test images. The data of participants M1 and M2 had their own peculiarities but also demonstrated a general increase of V-values with increasing eccentricity to 40-80° and decrease after reaching the maximum value.

One could propose various explanations of the described general shape of the curves and their inter-individual variability. However, it seems premature to do this now taking into account limited amount of the data obtained which is insufficient for thorough analysis although sufficient for certain definite conclusions. One of the evident conclusions is the necessity to assess the contribution of rods and ipRGCs to color perception since HSV color system appeared to be inappropriate for quantitative representation of the whole variety of the perceived colors.

## Discussion

Based on the obtained results, we conclude that the developed perimetric setup with two smartphones for displaying peripheral test and central reference stimuli proved to be suitable for comparison and matching in appearance color stimuli presented peripherally and centrally.

We would like to outline that, in the pilot series of experiments reported here, not all commonly employed means were used that otherwise would have allowed to increase precision of the measurements. In particular, in an attempt to reduce discomfort and fatigue usually experienced by observers in a typical study of peripheral vision, we did not apply bite bars, pupil dilatation, the Maxwellian view, or prolonged many-stage procedures worrying the participants. (Cf. a description of inconveniences caused by maintaining the state of the Maxwellian view during experiments in Neitz, (1990)).

Furthermore, we used a modified ACM procedure in that to ease the adjustment of the central reference stimulus, we forgone its flickering: thus, the peripheral test stimulus was turning on/off each 1.5 s while the central reference stimulus was presented without flickering. To ensure that this modification of the experimental procedure did not affect outcomes of the ACM, we conducted an experiment twice with three participant, i.e., with and without flickering of the central reference stimulus. As Figure S2 (Supplement) shows, the results from the two conditions did not differ significantly.

In spite of the simplifications and modifications mentioned above, the resulting accuracy of the measurements appears satisfactory to allow certain conclusions. Our data provide evidence that the proposed ACM technique can be employed over the range of eccentricities up to 95°, i.e. to the extreme VF (and retinal) periphery. Thus, we extended the findings of the psychophysical studies showing that deterioration of color vision with increasing eccentricity can be compensated by increasing the size and/or luminosity of the peripheral test stimuli (Noorlander et al., 1983; van Esch et al., 1984; Abramov et al., 1991; Murray et al., 2006; Tyler, 2015, 2016; etc.). Further, we present a quantitative description of inter-individual differences in color perception in the whole VF. Obtaining measurements in the wide range of eccentricities and comparing performance of several observers enabled us to get insight into causes of certain particularities and seemingly contradictory outcomes on the eccentricity-dependent color appearance reported previously, namely on constancy vs. changes of the perceived hue; enhancement vs. diminishing of saturation; the phenomena of supersaturation, etc. (Stabell & Stabell, 1976; Abramov & Gordon, 1977; Murray et al., 2012; Opper et al., 2014).

We are cognizant of limitations of the proposed technique, however, we hope that its main advantages – simplicity, affordability, and accessibility – could attract the attention of behavioral scientists and encourage the willing researchers to contribute investigation of appearance of color objects in the VF periphery.

Although the literature on color vision is huge, the majority of books and articles address the central vision only. The studies on peripheral color vision are scarce; many of these were carried out after a long training of the observers (e.g., Abramov & Gordon, 1977; Neitz, 1990). In many cases, only the authors could meet the challenges, i.e., master the experimental procedures and endure the experiment duration (e.g., Moreland & Cruz, 1959; Stabell & Stabell, 1979, 1982a,b, 1984, 1996; Nerger et al., 1995; Opper et al., 2014).

In comparison, in our experiments, most of the observers were able to perform all required measurements after a brief explanation and several short trials. This is encouraging for the data collection process: the duration of a smartphone ACM experiment comparable to that of sophisticated traditional experiments will open a possibility of collecting a larger database to address the peripheral vision phenomena in a greater scope.

Regarding the employment of smartphones and miniature computer devices for color vision investigation, recent literature shows a positive tendency both in laboratory research and in elaboration of color vision tests in field studies (see Dain et al., 2016; Bodduluri et al., 2017, 2018; Barbur et al., 2020; Pyayt, 2020). It is well known that one of the barriers to effective application of such devices is significant variability of colorimetric characteristics between devices, as well as instability of colorimetric parameters, which necessitates proper selection and calibration of the devices. Luckily, in this respect, the situation is improving quite rapidly: as a rule, each new model demonstrates a better colorimetric quality.

Finally, it is necessary to mention some important theoretical aspects of color vision study at the periphery of the VF projecting onto the periphery of the retina. The structural heterogeneity of the retina, addressed in the Introduction, apparently is reflected in its functional heterogeneity. It seems reasonable to suppose that retinal regions of different eccentricity are designed for both common functions and location-specific functions. This implies that comparing performance in the same visual task using the central and peripheral vision one could potentially reveal, if such relationships exist, functional similarity of the center and the periphery, or functional superiority of one over the other. For instance, it is known that the foveal area with the highest density of cone photoreceptors serves fine analysis of small stationary objects (that cannot be discriminated or detected at the periphery), whereas the peripheral vision serves detection of the large moving objects that enter the VF from its edge. In spite of this morphological and functional heterogeneity, certain visual tasks require functional involvement of the whole VF, in particular, color identification.

An expectation is that one can reveal either equivalence or superiority of either central or peripheral mechanisms depending on the particular visual task. Each of those tasks requires specific experimental paradigms and procedures. Due to the recent arrival of novel data on the contribution of rods to chromatic discrimination (e.g., Trezona, 1970; Buck, 2001; Cao et al., 2008) and ipRGCs to color perception (e.g., Cao & Barrionuevo, 2015; Schroeder et al., 2018; Zele et al., 2018; Spitschan, 2019), it becomes apparent that, instead of conventional 3-dimensional or 4-dimensional color spaces (Trezona, 1970, 1973, 1974; Brill, 1990; Smith & Pokorny, 2003; Polymeropoulos et al., 2011), a more appropriate model would require a 5-dimensional color space to describe full variety of colors perceived under certain experimental conditions (ambient light level, range of eccentricities, stimulus parameters). Moreover, an investigation of various neuronal mechanisms of color perception that combine inputs from the four photoreceptor types and ipRGCs could require fundamentally more complicated models than those developed by now. For example, even in 2007 Solomon and Lennie in their review “The machinery of colour vision” wrote: “…modern work has drawn attention to unexpected complexities of early organization … We describe, in the retina and in the lateral geniculate nucleus, many more pathways for colour signals than seemed possible only 15 years ago.” (Solomon & Lennie, 2007, p. 276).

It is, however, premature to expect rapid success in development of an adequate general model of the human peripheral color vision. The main barrier in this way is a substantial lack of information on both the machinery underlying the visual mechanisms (fine morphology and neurophysiology of the retina) and behavioral responses (psychophysical data) characterizing color perception. The complexity and heterogeneity of the human retinal structure and functions are still far from being studied in all details, neither are fully understood. Most psychophysical data have been obtained in specific artificial experimental conditions of eye fixation suppressing the peripheral mechanisms, which makes it problematic to puzzle out a general scheme of the peripheral color vision. In our view, further development of psychophysical experiments would require a greater variability of designs that would be more easily put into practice and enable capturing color vision phenomena under natural-viewing, ecologically valid conditions. Among various contemporary methods that are promising in this respect, we consider a technique of simulating central scotoma by means of contact lens (Almutleb et al., 2018; Rozhkova et al., 2019; Iomdina et al., 2020) and other accessible methods akin to the method described in the present paper. We are aware, though, that as relatively immature, such methods require further assessment and fine-tuning.

## Acknowledgement

We are greatly indebted to Prof. Galina Paramei for critical reading of our manuscript and helpful discussions, and to our colleagues, Prof. Nadezhda Vasilyeva and Anna Kazakova for their assistance in conducting our experiments.

## Disclosures

The authors declare no conflicts of interest.

## Financial support

Research was partially funded by Russian Foundation for Basic Research (19-015-00396A).

## Notes

### Competing Interest Statement

The authors have declared no competing interest.

