## Supplement for "A simple method for comparing peripheral and central color vision by means of two smartphones"

### *The effect of temperature on luminance of the primaries*

Since both luminance and spectral characteristics of displays are known to depend on temperature, it was necessary to assess corresponding dependencies for the smartphones employed. These characteristics are presented in Figure S1.

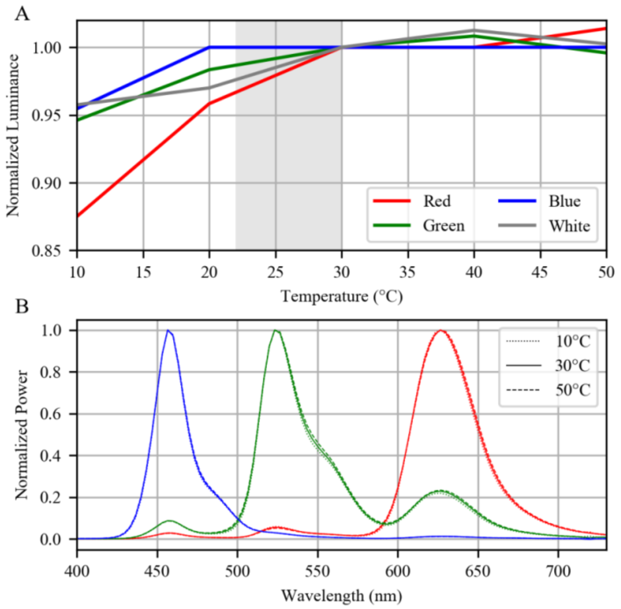


**Figure S1.** (A) The effect of temperature on luminance of white stimulus (gray line) and luminance of each primary (colored lines). Gray area indicates the range of temperature variation that is acceptable during matching experiments at room temperature of +22°C. (B) Spectral characteristics of the smartphone RGB primaries obtained at different temperatures.

The values of temperature indicated in Figure S1 correspond to the smartphone screen. This temperature was estimated with Fluke TiS60 thermal imager. To provide temperature variations we put smartphone in refrigerator cooling it down to +10°C or heat the smartphone with household hair dryer up to +50°C. In the case of usual laboratory temperature of +22°C, the static stimulus spot of maximal brightness achieved stationary temperature of about +30°C during 2-3 minutes, flickering spot achieved stationary temperature of +26°C. Luminance variations of the R, G, and B primaries were less than 5% in the temperature range from +20°C to 50°C (Figure S1A).

The spectral variations of the R, G, B primaries in the temperature range from +20°C to 50°C could be considered as insignificant (Figure S1B).

### *The effect of test stimulus flickering on results of the center-periphery color matching*

To ease the adjustment of the central reference stimulus we forgone its flickering, while the peripheral test stimulus was flickering (1.5 s “on” + 1.5 s “off”). We checked that such modification of the experimental procedure did not influence significantly the results of the ACM. It is illustrated in Figure S2 for the case of V-estimates.

For obtaining the relationship between the reference image brightness parameter V in the HSV color model (varied from 0 to 1) and the test image luminance $L_{sm}$ (expressed in 8-bit 256 RGB values) in the software for image generation, two smartphones were placed side by side and the participants were allowed to shift the gaze from one to the other for matching the images. Figure S2 shows the results of such matching for the case of the stationary test and reference images (solid lines) and for the case when the test image was turning on/off each 1.5 s (dashed lines).

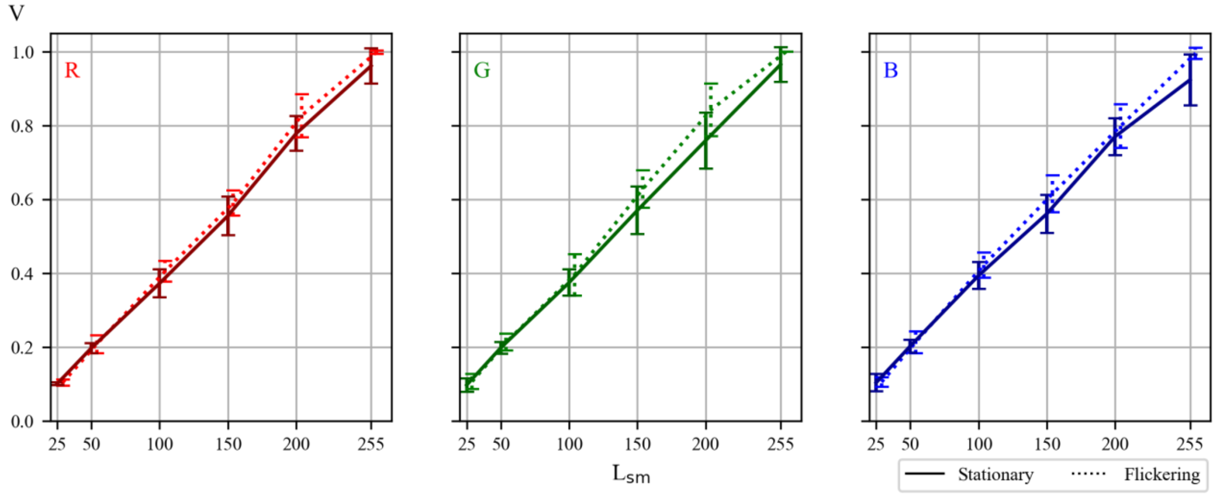


**Figure S2.** Comparison of the results obtained in conditions of matching stationary reference stimuli to stationary (solid line) and flickering (dashed lines) test stimuli. Each point represents the average of 15 measurements (3 participants, x 5 trials). Error bars represent ±SEM.

Comparison of the results obtained with stationary and flickering test images was necessary to be sure that our standard stimulus duration of 1.5 s was enough to mimic stationary test stimulus and that the inter-stimulus interval of 1.5 s was sufficient for recovery. The data presented in Figure S2 could be used to assess the reliability of the main results of our investigation. The curves obtained for the average V-estimates in the cases of flickering and stationary stimuli and corresponding standard errors of the mean values determine the accuracy of all V-estimates indicated in the main text body.

### *The effect of interchanging the test and reference smartphones on the estimates of H, S and V*

In order to assess the effect of small differences in individual spectral characteristics of smartphones on color marching results, we compared the values of HSV-estimates in conditions of interchanging the test and reference smartphones of the pair Samsung Galaxy S8. Figure S3 shows the data of two participants obtained in three experimental sessions: with certain initial positions of the test and reference smartphones (1), after swapping the smartphone positions (2), and after returning the smartphones into their original positions (3). It is noteworthy that

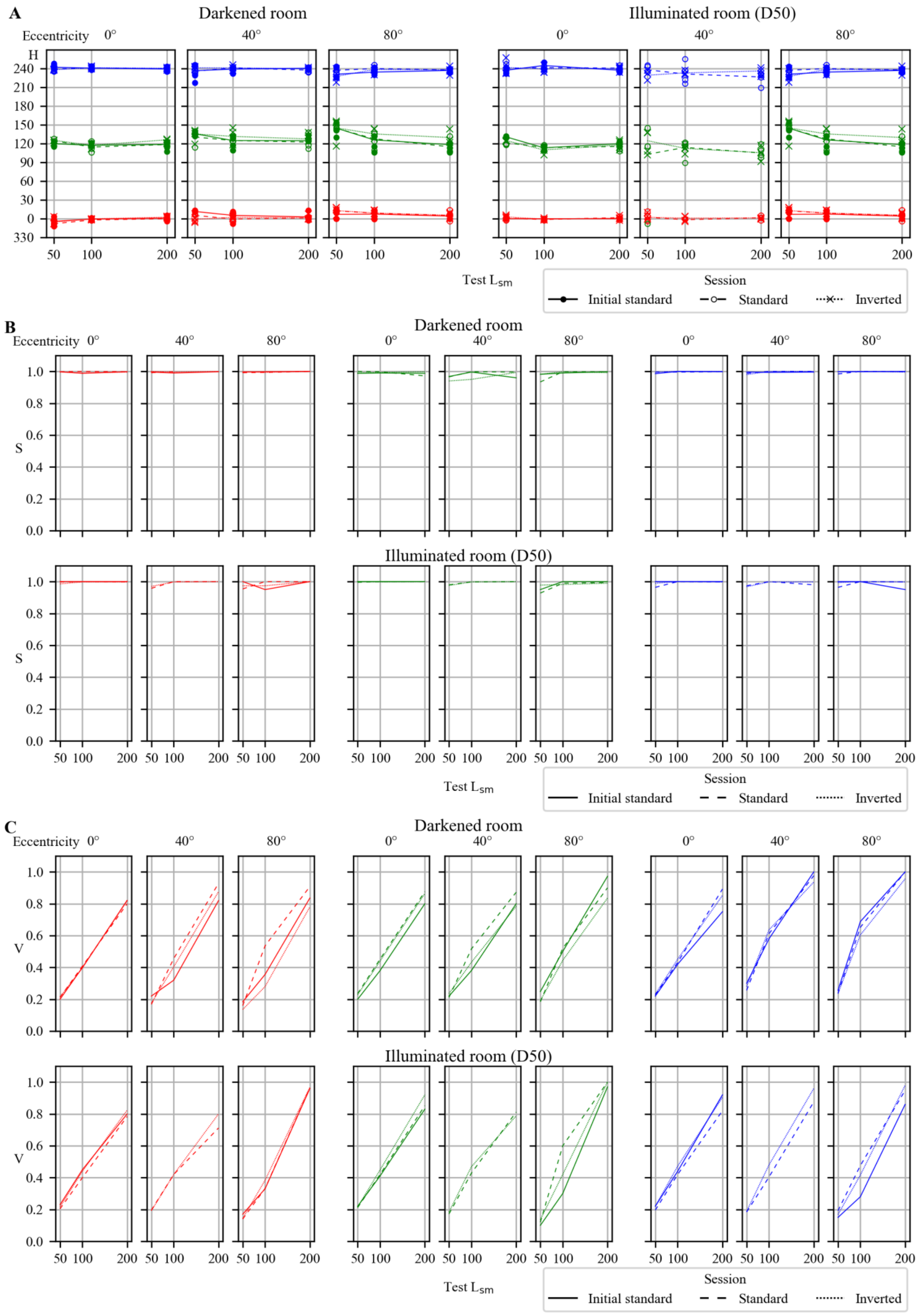
A
Darkened room
Illuminated room (D50)
Eccentricity 0°
40°
80°
H
240
210
180
150
120
90
60
30
0
330
50 100 200
Test $L_{sm}$
Session
Initial standard
Standard
Inverted
B
Darkened room
Illuminated room (D50)
S
1.0 0.8 0.6 0.4 0.2 0.0
Test $L_{sm}$
Session
Initial standard
Standard
Inverted
C
Darkened room
Illuminated room (D50)
V
1.0 0.8 0.6 0.4 0.2 0.0
Test $L_{sm}$
Session
Initial standard
Standard
Inverted

**Figure S3.** The data obtained for perceived hue (A), saturation (B), and brightness (C) in one and the same observer in three experimental sessions: with certain initial positions of the test and reference smartphones (1), after swapping the smartphone positions (2), and after returning the smartphones into their original positions (3). The two experiments were performed under dark adaptation and in an illuminated room (see the headings over individual graphs); *x*-axis: test stimulus intensity in RGB-scale; *y*-axis: H-estimates (A), S-estimates (B), and V-estimates (C) in the HSV color coordinate system for the smartphone primaries R, G, and B.

As one could conclude from the Figure S3A, the anticipated influence of the test-reference spectral differences on H-estimates was noticeable but not much: in 15 from 18 sets, all three curves in each triad appeared to be quite similar indicating that the small differences in the individual spectral characteristics of the smartphones did not affect the results of matching significantly.
